# Electrical-charge accumulation enables integrative quality control during *B. subtilis* sporulation

**DOI:** 10.1101/349654

**Authors:** Teja Sirec, Pauline Buffard, Jordi Garcia-Ojalvo, Munehiro Asally

## Abstract

Quality control of offspring is important for the survival of cells. However, the mechanism by which quality of offspring cells may be monitored while running genetic programs of cellular differentiation remains largely unclear. Here we investigated a quality control system during *Bacillus subtilis* spore formation by combining single-cell time-lapse microscopy, molecular biology and mathematical modelling. Our results revealed that the quality-control system via premature germination is coupled with the accumulation of cations on the surface of developing forespores. Specifically, the forespores accumulating less cations on their surface are more likely to be aborted. This charge accumulation system enables the projection of multidimensional information about the external environment and morphological development of the forespore onto a one-dimensional information of cation accumulation. Based on the insight we gain, we propose a novel use of Nernstian chemicals for reducing the yield and quality of *Bacillus* endospores.

## INTRODUCTION

The significances of electrophysiological dynamics in bacteria has been uncovered in the last few years (1). Cells within *Bacillus subtilis* biofilms can communicate with each other through electrical signaling mediated by the gating of potassium channels (2). This bacterial electrical signaling increases the fitness of the population by enabling and metabolic co-dependence within a biofilm (3) and nutrient time-sharing of distant biofilms (4). A recent study also revealed that the bacterial electrical signaling can attract motile cells in a species-independent manner (5). In *E. coli*, rapid change in membrane potential mediated by the opening of calcium channels is crucial for the response to mechanical stresses (6). As a result of these pioneering studies, bacterial electrophysiology concerning the ion dynamics has become an exciting avenue of research, which is however still a largely uncharted field in microbiology. A particularly important unanswered question is how electrophysiological dynamics interplays with complex genetic processes (*e.g.* cellular differentiation). Notably, such interplay between biophysical states (*e.g.* membrane potential, redox states, cell size) and genetic processes is a timely research topic not only in bacterial electrophysiology but also in various research areas of quantitative biology and synthetic biology (7–10). With this in mind, we investigated the *B. subtilis* spore formation with a focus on the dynamics of cationic molecules and spore quality control.

Sporulation of *B. subtilis* is among the best-characterized bacterial cellular differentiation processes (11, 12). Over five decades of in-depth genetic studies and high-throughput analyses have identified the genes and proteins governing this differentiation process (13, 14). The principles of the cellular decision making, leading to the commitment to sporulation, have been deciphered by single-cell time-lapse microscopy and mathematical modelling (15, 16). These extensive body of research has resulted in a good understanding of the genetic regulation during sporulation. Briefly, the differentiation into endospores begins with reversible phosphorylation of the master transcription factor Spo0A via phosphorelay. The phosphorylation of Spo0A leads to the sporulation commitment defined by the irreversible formation of asymmetric septum. The commitment triggers the multistage differentiation program regulated by compartment-specific sporulation sigma factors (*σ^F^*, *σ^E^*, *σ^G^*, *σ^K^*), undergoing a series of genetically regulated developmental processes; *i.e.* forespore engulfment, cortex synthesis, coat assembly and mother-cell lysis.

While it is well established that sporulation is regulated by a complex genetic program, it is also evident that the process is subject to a variety of internal and external conditions (Figure 1A). Poorly developing spores are eliminated from the population through quality control systems mediated by Clp proteases and GerA-dependent premature germination (17, 18). Intriguingly, the quality (*e.g.* resistance and germination properties) of endospores differs depending on the environmental conditions in which sporulation takes place (19). For instance, endospores produced at high temperature (~50 °C) exhibit higher resistance against heat (~100 °C) (20). The thickness of spore coats varies depending on the culture medium (21). Strikingly, even with an identical medium and temperature, the endospores prepared in liquid are less resistant to heat than the endospores produced on plates (22). It is worth noting that such condition-dependent variability of spores is recognized as a major challenge in food industries since it precludes the reliable standardization of the sterilization procedures (23, 24). However, despite these observations demonstrating the phenotypic plasticity of endospores, the mechanism by which diverse environmental conditions (*e.g.* media compositions, temperature, pH) and morphological properties of developing spores (*e.g.* cortex synthesis, and coat assembly) influence the sporulation process remains unknown.

**Figure 1.**
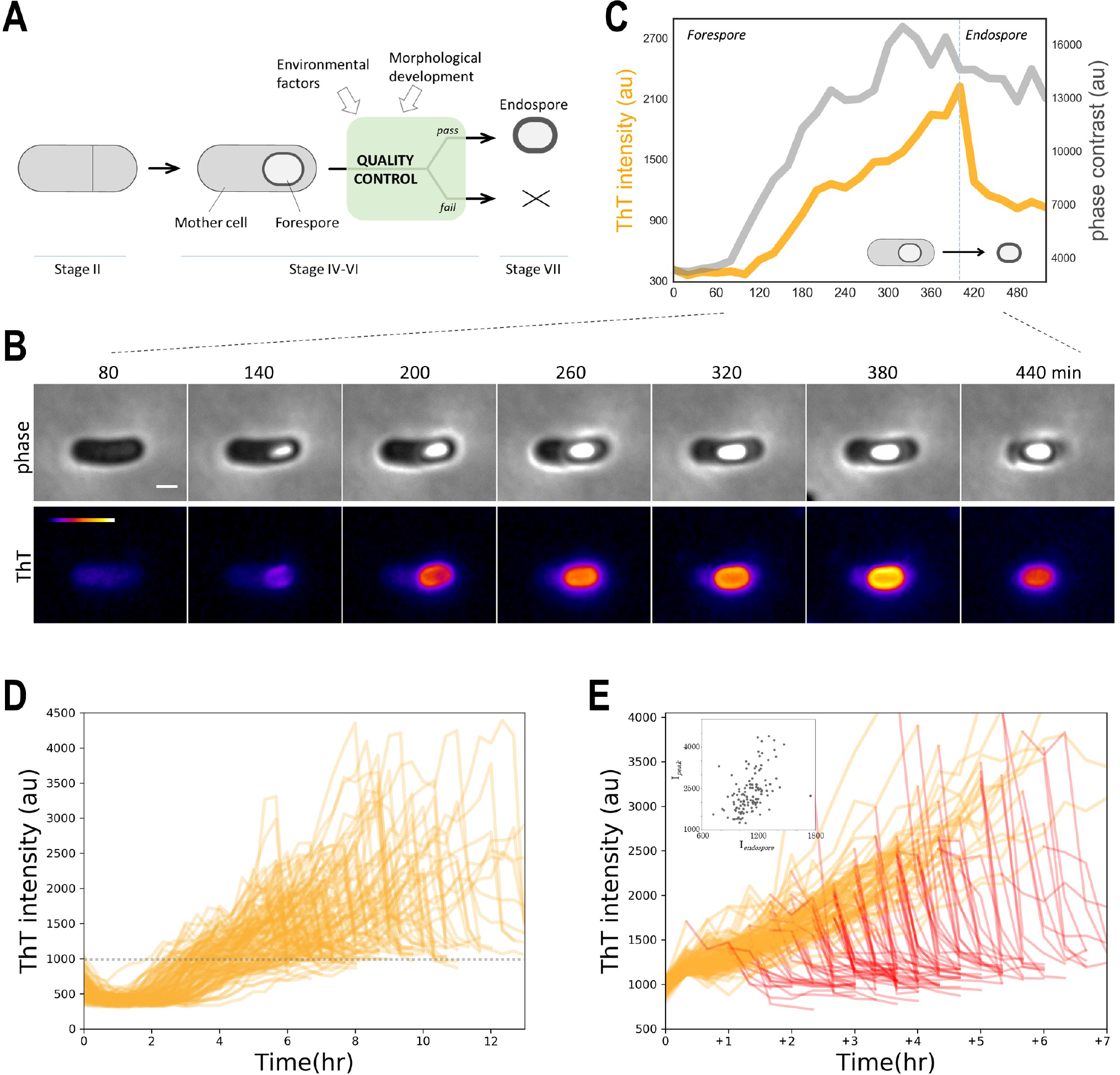
Single-cell dynamics of a cationic molecule during *B. subtilis* sporulation revealed a gradual increase and sudden drop upon mother-cell lysis. A) Illustrative diagram shows the sporulation stages. Asymmetrically divided cells (Stage II) undergo engulfment (Stage III), and late sporulation (cortex synthesis (Stage IV), coat assembly (Stage V), and maturation (Stage VI). During the late sporulation stages, cells accumulate cations. B) Film-strip images showing the phase-contrast channel (upper) and Thioflavin T (ThT) fluorescence channel (lower) of a live single-cell sporulating cell. The scale bar is 1 μm. Color scale for ThT intensity is shown in the left-end panel for ThT. Film-strip is a representative of successfully sporulating cells from twenty-nine independent experiments. C) Time series of the mean intensities of phase contrast channel (grey) and ThT fluorescence (orange) in forespore/endospore region. The data corresponding to the forespore region of the sporulating cell shown in panel B. Time series shows a gradual increase of ThT, which drops upon mother-cell lysis. D) Time series of single-cell ThT fluorescence dynamics on forespores (n = 122) from two independent experiments. Forespore regions of ThT intensity were measure until endospores are released from mother cells. All time-series qualitatively follow the pattern represented in panel C, but a great degree of heterogeneity between cells. Dashed line represents the value used for alignment in panel E. E) The data set shown in panel D is aligned to the frame it first reaches ThT intensity value 1,000 (shown with a dashed line in panel D). Time series after reaching maximum are highlighted in red. Time series show a relatively finite slope of ThT intensity increase and a great degree of heterogeneity in timing of mother-cell lysis (red lines). Inset plot shows the ThT intensities at peak and after released from mother cells for each time series. See also Figure S2.

We hypothesized that diverse morphological properties and environmental factors can be sensed integratively through electrophysiological dynamics of cells during sporulation, which provides an orthogonal dimension to the complex genetic differentiation program. We characterized the dynamics of cationic molecules during late sporulation (stage IV-VI) of *B. subtilis* at the single-cell level (Figure 1A). Our fluorescence time-lapse microscopy measurements, combined with examinations of genetic mutant strains and simple computational simulations, suggest that the success rate of endospore formation depends on the electrical attraction of cations to the forespore surfaces. More specifically, spore formation is more likely to be completed when forespores accumulate high levels of cations on their surface. We found that the electrical attraction during late sporulation provides a mechanistic explanation of the quality-control system mediated by premature germination. Intriguingly, this sensing mechanism mediated by the electrical attraction enables the quality-control system to be responsive not only to morphogenetic errors of forespores but also to the external environments during sporulation. Thus, this mechanism could provide an integrative monitoring system for the bacterial spore-quality-control mechanism. Finally, in the light of the insights we obtained, we succeed in promoting the failure of sporulation using a Nernstian chemical Thioflavin T (ThT), while at the same time decreasing the resistance property of endospores against wet-heat.

## RESULTS

### Single-cell measurements of membrane potential dynamics during sporulation

To characterize the ion dynamics during sporulation and its potential role in quality control, we began by measuring the membrane potential dynamics in live sporulating *B. subtilis* cells using the positively charged Nernstian dye ThT (2, 4, 5). We specifically focused on the late sporulation stages because herein the quality-control systems operate (Figure 1A) and the cells are known to accumulate a high level of monovalent and divalent cations (25). Our fluorescence measurements of ThT revealed a gradual increase of the signal on a developing forespore, which then rapidly drops upon mother-cell lysis (Figures 1B and 1C, and Movie 1). The fluorescence signal was seen intensely on the peripheral regions of the forespores (Figure 1B). The increase in fluorescence intensity is preceded by the increase in forespore brightness in phase-contrast channel (Figure 1C). We also measured the pH dynamics during sporulation using the ratio-metric pH sensor pHluorin. Consistent with previous studies performing population-level measurements (26), our single-cell imaging showed that the pH inside the core compartment decreases dramatically during late sporulation (Figures S1A and S1B, and Movie 2). The acidification of the core region appeared to coincide with the increase in phase brightness, suggesting that acidification of the core occurs before the increase in ThT intensity on the forespore periphery. The rapid drop of the ThT signal upon mother-cell lysis suggests that the mother cells accumulate this cationic dye during sporulation by dynamic association.

Although ThT has been extensively used with *B. subtilis* as the indicator for membrane potential (2, 4, 5), it is also a well-known fluorescent marker for amyloid fibrils. It has been proposed that endospores may have amyloid fibrils on their surfaces which can be stained by ThT (27). The rapid drop of ThT signal upon mother-cell lysis (Figures 1B and 1C) suggests that ThT is not reporting amyloid fibrils since amyloids fibrils are stable and unlikely to depolarize in such time scale. However, there remained a possibility that ThT is reporting the formation of amyloid-like structures on forespore surfaces. To examine whether ThT is faithfully reporting the electrical interactions in our experiments, we used another fluorescent indicator for membrane potential, namely, Tetramethylrhodamine methyl ester (TMRM). TMRM is a positively charged lipophilic potentiometric dye that accumulates electrophoretically, but it does not bind amyloid fibrils. Fluorescence time-lapse microscopy showed that the dynamics of TMRM and ThT signals during sporulation are indistinguishable (Figures S1C and S1D, and Movie 3). Therefore, we concluded that these fluorescent markers are indeed reporting the increase of the electrophoretic attraction on the forespore surface. Together, our data show that the forespore surface gradually becomes electrically negative, which attracts positively charged molecules. This interpretation agrees with the literature reporting the negative surface potential of fully developed endospores (zeta potential −26 mV at pH 7.0) (28).

We next performed single-cell tracking of ThT dynamics with the cells that produced phase-bright endospores (*n* = 122). For individual cell, ThT intensity on the forespore compartment was measured until 1 hour after the lysis of mother cell. All time-series from successfully sporulating cells follow the pattern presented in Figures 1B and 1C; namely, ThT intensity increases gradually and drops rapidly upon mother-cell lysis. However, there is a large degree of heterogeneity among individual cells (Figure 1D). It is known that the late sporulation in *B. subtilis* is highly heterogeneous, which has made the interpretation of population-level measurements challenging. To better understand the sources of heterogeneity, we analyzed the ThT time-traces of individual cells. Specifically, we aligned each of the time-trace to the frame where the ThT intensity reaches 1,000 AU (arbitrary unit), shown as a dashed horizontal grey line in Figure 1D. This alignment revealed a consistent increase rate of ThT signal among cells (Figure 1E). On the contrary, the length of time it takes to reach mother-cell lysis varies substantially among cells (Figure 1E; the time-traces after mother-cell lysis are highlighted in red). On average, after the ThT value reaches 1,000 AU, it takes 3.28 hours for the spores to be released from mother cells with a standard deviation of 1.44 hours (Figure S2). As a consequence of the finite gradual increase and heterogeneous timing of ‘exit’, different cells reach the different level of peak ThT intensities; the longer the forespore persists in the mother cells, the greater the ThT accumulation becomes on forespores. However, the peak ThT intensity levels are only weakly correlated with the intensity levels on the released endospores (Figure 1E, inset). This result suggests that not only the negative surface potential of fore/endo-spores but the internal environment of mother cells may also contribute to the accumulation of ThT on forespore periphery. In summary, our single-cell experiments revealed the dynamics of negative surface potential of forespores. We showed that ThT (a positively charged dye) is increasingly accumulated on the forespore surface as the surface becomes negative.

### Outer spore coat accumulates positive molecules on the forespore surfaces by electrical attraction

We next aimed to understand the sources of negative surface potential build-up during late sporulation. The negative surface potential of developed endospores is known to be implicated with the spore-coat layers (29). Therefore, we hypothesized that the attraction of positively charged dyes to forespore surfaces is also due to the outer coat layers. To examine this conjecture, we measured the ThT dynamics with mutant strains lacking the outer spore coat; specifically, we utilized two deletion mutant strains lacking *sigK* and *gerE*. These genes encode late sporulation regulators which control the expression of coat proteins and cortex synthesis enzymes. The fluorescence intensities on forespores in these mutant strains were clearly lower than in the wildtype strain (Figures 2A and 2B, see also Figure S1D). These results suggest that the outer coat layer is a main contributor to the accumulation of positive ions on the forespore surfaces. To further examine the contribution of individual coat proteins, we measured ThT fluorescence with the mutants lacking the main structural proteins of the outer spore coat. Specifically, we measured ThT intensities on endospore surfaces with the mutant strains of *cotB, cotBG, cotC*, and *cotCU*, as well as the *gerE* strain. The mutants mostly showed similar intensity level of ThT compared to wildtype, however, a slight decrease in ThT intensity was observed with the *cotBG* double deletion strain (Figure 2C). Therefore, the negative surface potential of forespores is likely associated with the multiple components of outer endospores coat, including CotB and CotG. Together with previous studies reporting the negative electric charge of endospores (28), these results suggest that the finite gradual increase of Nernstian fluorescence signal (Figure 1E) is reflecting the assembly of the coat layers, one of the main protective layers of spores.

**Figure 2.**
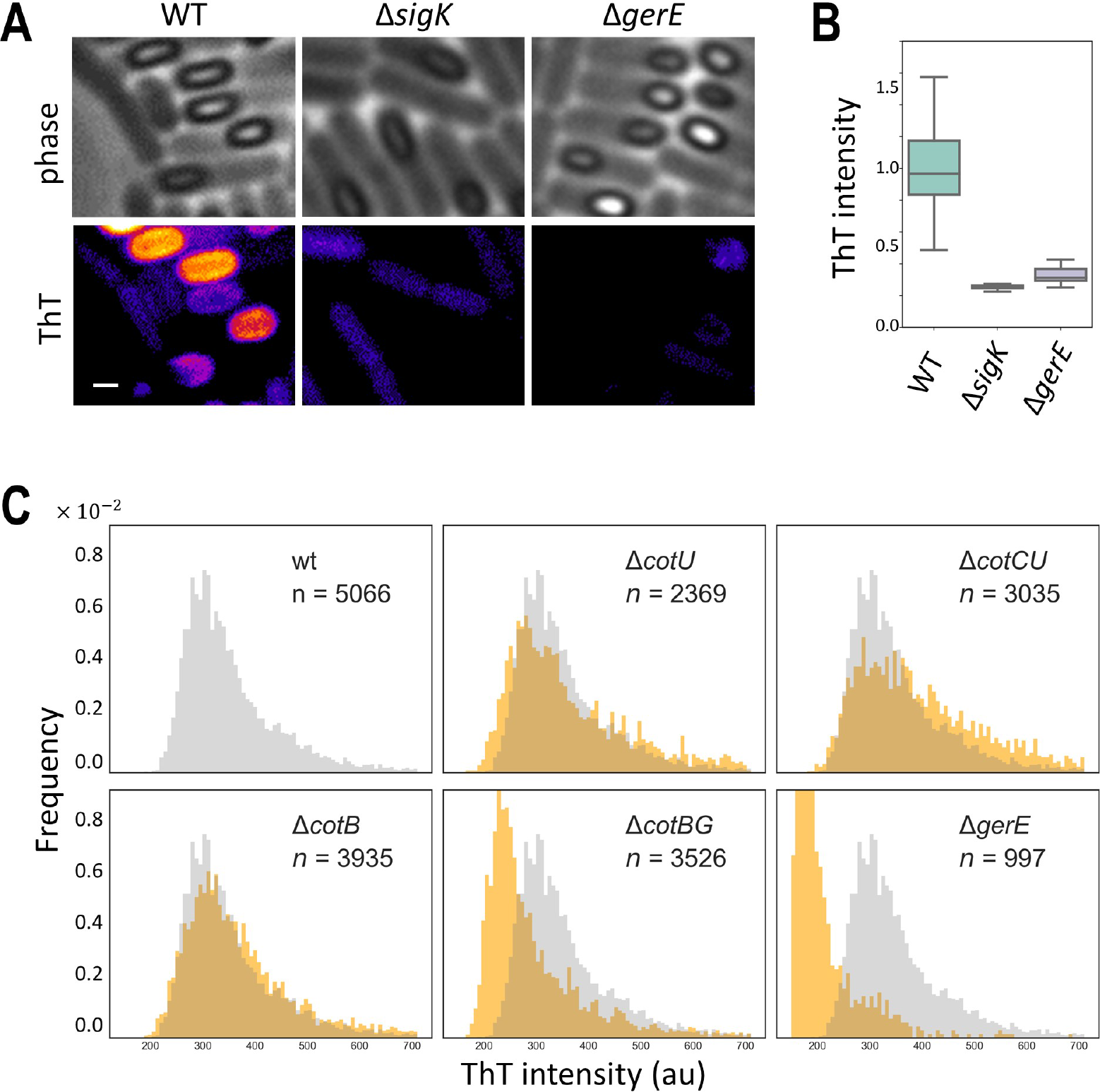
ThT is accumulated on forespore surfaces through electrical attraction by spore protective layer. A) Microscopy images of phase contrast (upper panels) and ThT fluorescence (lower panels) of wildtype (wt), *ΔsigK*, and *ΔgerE* strains. Scale bar, 1 μm. B) Box and whisker plot showing peak ThT intensities in wt, *ΔsigK*, and *ΔgerE* strains (*n* = 30). The boxes and whiskers show IQR (interquartile range), and 1.5× IQR, respectively. This plot shows weaker ThT signal with the *ΔsigK*, and *ΔgerE* strains. C) Histogram of ThT intensities on endospores with wt, *ΔcotU, ΔcotCU, ΔcotB, ΔcotBG*, and *ΔgerE*. The number of endospores analyzed is indicated in each panel. Histogram of wt strain is shown in grey in all panels for comparison with mutant strains (shown in orange). *ΔcotBG*, and *ΔgerE* strains exhibit slightly lower ThT intensity levels.

### Electrical attraction of cations on forespore surfaces correlates with premature germination frequency

We next wondered if the cation accumulation on the surface of forespores has a biological role in sporulation. Through single-cell time-lapse microscopy, we noticed that some phase-bright forespores turned into phase-dark while still inside mother cells (Figure 3A; upper panel, Figure 3B, and Movie 4). This change in the phase contrast was reminiscent to the germination of endospores (30). We thus speculated that the change in the phase-contrast is due to premature germination within mother cells. To test this idea, we conducted time-lapse microscopy with the mutant strain lacking *gerA* gene, which encodes a main germinant receptor essential for the L-alanine-induced germination. As expected, the deletion of *gerA* gene eliminates the change of phase brightness with forespores (Figure 3A; lower panel, Figure 3B, and Movie 5). This observation appeared complementary to the genetic analyses of a recent paper (18). This study by Ramirez-Guadiana *et al* elegantly demonstrated that GerA-mediated premature germination can act as a quality-control system of forespore morphogenesis; *i.e*. forespores germinate prematurely when there are morphogenetic errors. However, how cells are able to couple premature germination with a range of qualitatively different types of morphogenetic errors, such as synthesis of the endospore protective layers (coat and cortex) and core dehydration, remained unclear. We thus investigated the possible mechanisms by which the frequency of premature germination is tuned during sporulation.

**Figure 3.**
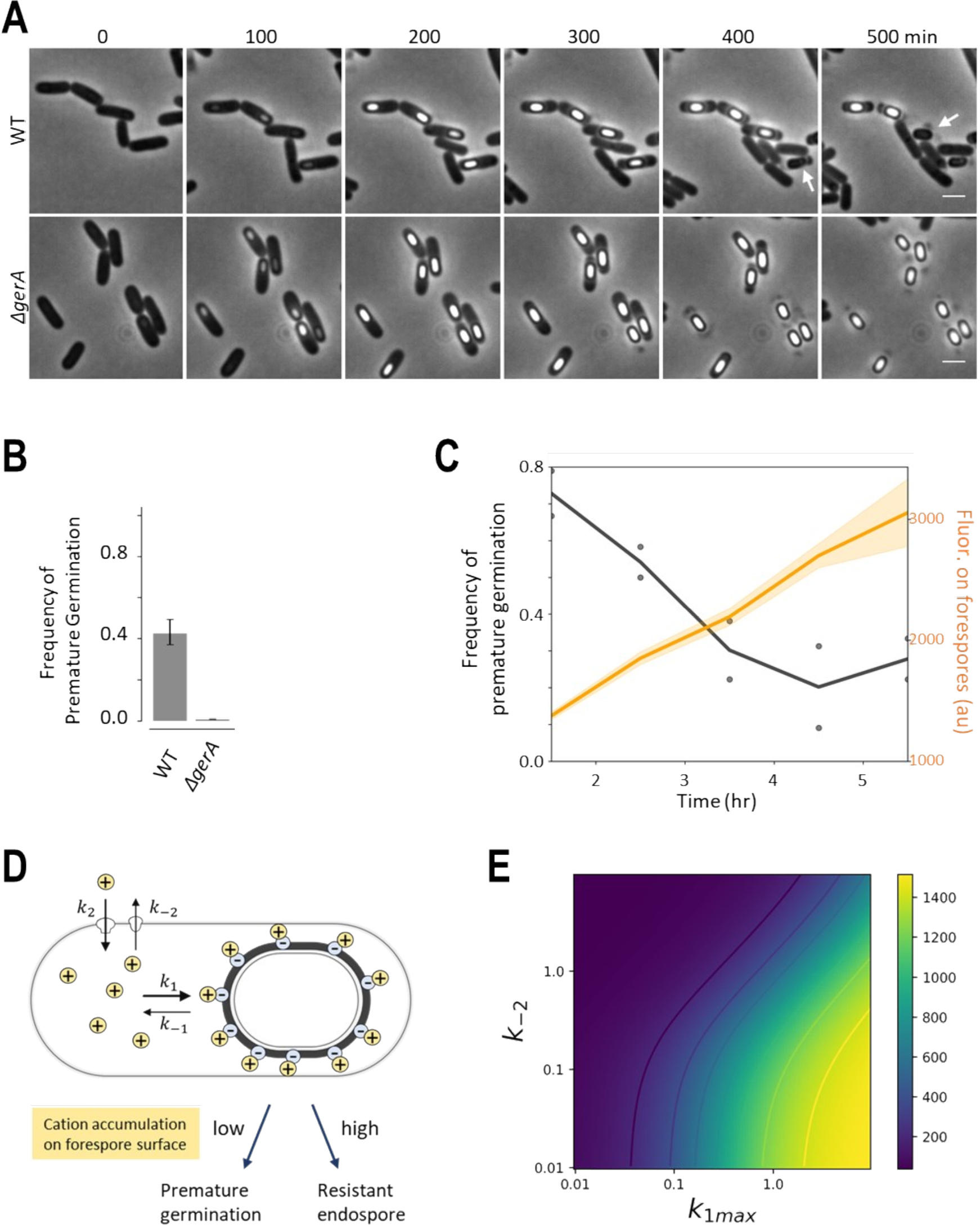
Accumulation levels of cations on forespore surface inversely correlate with the frequency of premature germination. A) Film-strip images of wildtype (WT) and *ΔgerA* strains during sporulation in the phase-contrast channel. Scale bar, 2 μm. Several wildtype forespores undergo premature germination (indicated by white arrows), while all forespores complete sporulation in *ΔgerA*. B) Quantification of frequency of premature germination in wt and *ΔgerA* strains (at least 351 sporulating cells were analysed for each strain/condition). Error bars are 95 % confidence intervals for Poisson distribution. C) Frequency of premature germination decreases as ThT signal on the forespores increases. Time series of single cell ThT intensity are aligned as Figure 1E and frequency of premature germination was calculated for each hour. D) A diagram shows the hypothesis that premature germination frequency is coupled with the accumulation levels of cations on forespore surfaces. A simple mathematical model was developed accounting the influx and efflux of cations (*k*_1_ and *k*_−1_) and the binding and unbinding of cations to forespore surfaces (*k*_2_ and *k*_−2_). Accounting for the increase of surface negative potential of forespores, we assume *k*_1_ grows as a function of time with the equation: *k*_1_(t) = *k*_1*max*_*t*/(*t_m_*+*t*). E) Numerical simulation of the model showed monotonic decay of cation accumulation on forespore surfaces (*C_f_*) by decreasing *k_1max_* (x-axis) or increasing *k*_−2_ (y-axis). The simulation was performed with the following parameter settng; *t_m_* =1, *k_2_* = 25, *E* = 5, k_−1_=0.1.

Taking an advantage of time-lapse single-cell imaging, we quantified the time evolution of premature germination frequency. Our data revealed that the frequency of premature germination decreases as the level of cation accumulation on the surface of forespores increases (Figure 3C). Therefore, we hypothesized that the accumulation of cationic ions on forespore surfaces may prevent the germination of forespores (Figure 3D).

Intriguingly, it has been shown that high levels of various cationic ions (*e.g*. K^+^, Na^+^ and Ca^2+^) limit the access of L-alanine to the intermembrane space where GerA exists, and consequently prevent the germination of endospores (31). Therefore, the premature germination of forespores may also be affected by cations in a manner similar to the germination of fully developed endospores. To better understand which native cations may be involved in the modulation of premature germination frequency, we used fluorescent indicators APG-2 AM (Asante Potassium Green-2 Acetoxymethyl), ANG-2 AM (Asante Natrium Green-2 Acetoxymethyl) and Fluo-4FF AM to measure potassium, sodium, and calcium, respectively. Potassium and sodium appeared to be accumulated on forespore periphery regions, while calcium is localized in the spore core (Figure S4A). APG-2 provides a clear signal, while the other two reporters are relatively noisy with our experimental setting. Therefore, although obviously we do not exclude the possibilities that other cations (*e.g*. manganese) are also accumulated, we concluded that potassium is one of the cations that accumulates highly on the forespore surfaces. The quantification of APG-2 signal showed that the potassium levels on forespores are slightly lower with presence of ThT in the media, suggesting that ThT may act as a competitive inhibitor for potassium in terms of the accumulation to forespore surfaces (Figure S4B).

To investigate the hypothesis that the potassium accumulation on forespore surfaces suppresses the frequency of premature germination, we established a simple mathematical model describing the dynamics of the cations in cytoplasm (*C_c_*) and forespore-bound (*C_f_*) (Figure 3D). Accounting for the development of negative surface potential of forespores, we assumed that the affinity of cations to forespore surfaces (*k*_1_) increases over time which eventually saturates at a value, *k*_1*max*_. Numerical simulations of the model showed that *C_f_* decreases in a monotonic manner when cation accumulation is inhibited (lower *k*_1*max*_) or cation efflux is increased (higher *k*_−2_) (Figure 3E).

### The premature germination frequency can be modulated by chemical perturbations

We first examined the prediction from the model regarding the decrease in cation accumulation (lower *k*_1*max*_). Taking into account the charge conservation law, it is expected that exogenous cations should act as a competitive inhibitor for native cations (*e.g*. potassium) in terms of forespore surface accumulation. In fact, we observed decreased accumulation of potassium on forespore surfaces when ThT is present (Figure S4B). Therefore, our model suggested that addition of ThT should slow the growth of *k*_1_ and thus increase the premature germination frequency. We measured the frequency of premature germination with and without ThT, and found that a greater fraction of forespores, albeit only slightly, germinate prematurely when ThT is supplemented to the media (Figure 4A). We confirmed that the premature germination remains absent in Δ*gerA* strain with or without the presence of ThT (Figure 4A). We also examined whether ThT has a direct interaction with the GerA receptor, by conducting germination assay with purified endospores. The result showed that, unlike premature germination of forespores, the germination of mature endospores is unaffected by ThT (Figure S3). This result indicates that observed increase of premature germination frequency is not due to direct interaction between GerA receptor and ThT. To examine the causality of the ThT effect, we tested different ThT concentrations and measured the premature germination frequency. The results showed a clear increase of phase-dark endospore fraction as a function of ThT concentrations (Figure 4B and 4C).

**Figure 4.**
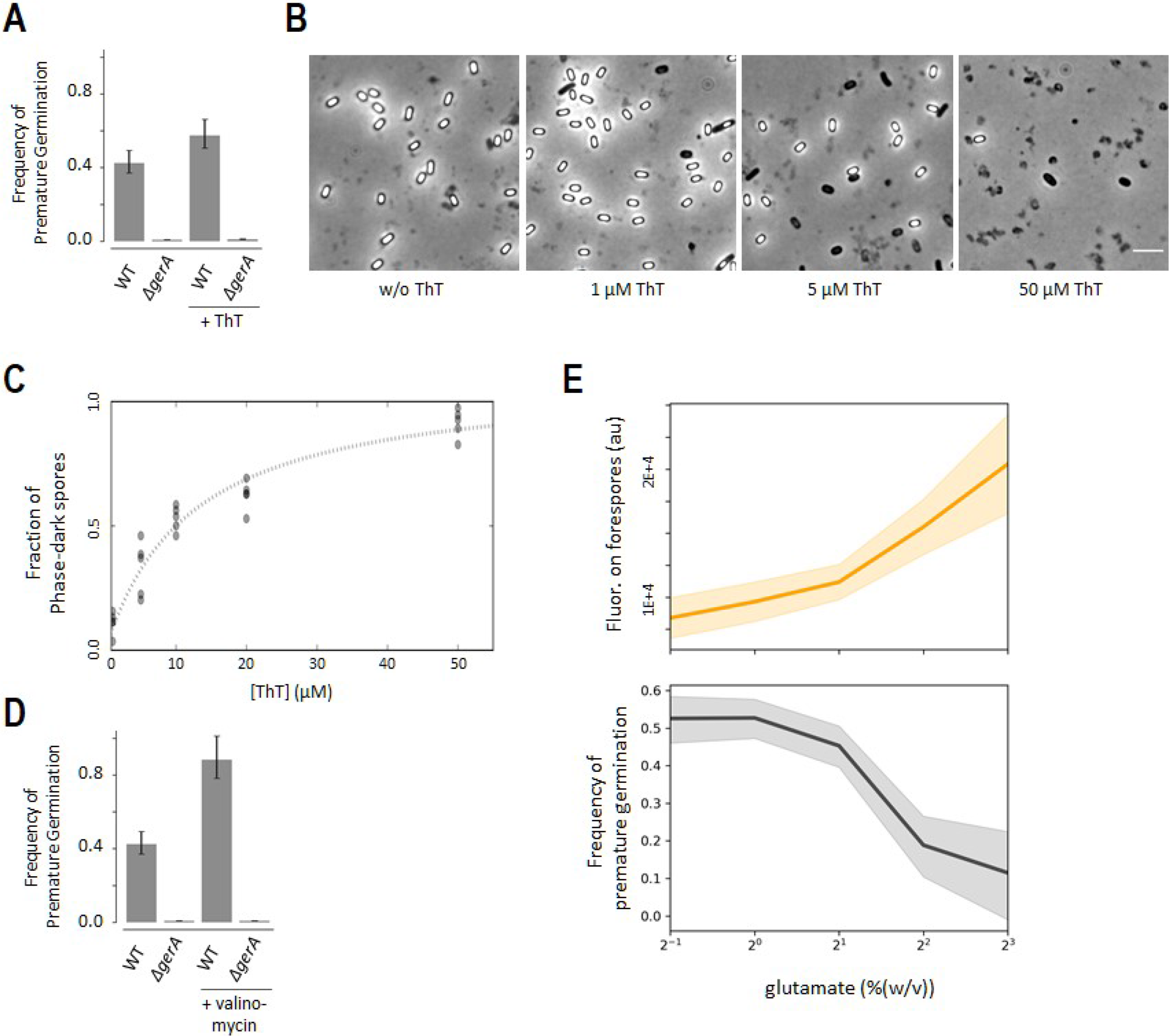
The frequency of premature germination can be modulated. A) Sigle-cell microscopy analysis revealed that the frequency of premature germination is slightly higher when the media is supplemented with 10 μM ThT. The premature germination remains absent with gerA deletion strain. At least 351 sporulating cells were analyzed for each condition/strain. The error bars are 95% confidence intervals for Poisson distribution. B) Microscopy images of endospores cultured in liquid RM supplemented without or with ThT at 1 μM, 5 μM, 50 μM. Scale bar, 5 μM. C) A plot showing the fraction of germinated (phase-dark) endospores. More endospores are phase-dark with higher ThT concentration supplemented to the media. Each dot represents experimental result. D) Addition of valinomycin to the media increases the frequency of premature germination. At least 339 sporulating cells were analyzed for each strain/condition. Error bars represent 95% confidence intervals for Poisson distribution. Sporulation of *gerA* strain appeared unaffected by valinomycin. Corresponding microscopy images in Figure S5. E) The peak fluorescence signal on forespore surfaces (TMRM) increases as a function of glutamate levels in the media (upper plot), while the frequency of premature germination drops (lower plot). Shaded regions are standard deviation for the upper plot, and Poisson confidence intervals for the lower plot.

We next examined the prediction from the model regarding the increase in cation efflux (higher *k*_−2_) by utilizing valinomycin. Valinomycin is a potassium ionophore, meaning it increases the membrane permeability of potassium, thus elevates *k*_−2_ (potassium efflux). As can be seen in Figure 3E, the model predicts that valinomycin should increase the frequency of premature germination. In order to minimize the potential global impacts of valinomycin, we exposed cells to valinomycin only after cells reach the commitment stage (cultured in RM for 3.5 hours). Quantification of single-cell time-lapse microscopy data showed that the frequency of premature germination significantly increases by valinomycin (Figure 4D). To check that this is not due to a general toxic effect of valinomycin, committed cells of Δ*gerA* strain were exposed to valinomycin. Contrary to the results with wildtype, the sporulation of Δ*gerA* strain was unaffected by valinomycin and phase-bright endospores were produced normally (Figures 4D and S5). This result indicates that the crucial role of the transmembrane electrochemical gradient of potassium during late sporulation (stage IV-VI) is to prevent premature germination, but not to support harvesting the energy required for the completion of late sporulation process.

### Glutamate availability influences the frequency of premature germination

While a previous study suggested that the frequency of premature germination may be constant in different media (18), our model predicted that environmental conditions, that affect the membrane potential, should alter the frequency of premature germination. To this end, we focused on glutamate availability because glutamate is a gating molecule for the *B. subtilis* potassium channels which remain closed in glutamate rich media (2, 4). Thus, according to Figure 3E, our model predicted that premature germination frequency would decrease as a function of glutamate levels in the media (lower *k*_−2_).

To test this experimentally, cells committed to sporulation (cultured in RM for 3.5 hours) were transferred to RM containing different levels of glutamate (0.5 - 8.0 % (w/v)). The frequency of premature germination and the forespore-surface charge were determined by single-cell microscopy (Figure 3B). The results showed that the frequency of premature germination decreases as a function of glutamate concentrations, while the accumulation of charge is inversely correlated with the glutamate levels (Figure 4E). This result indicates that the frequency of premature germination is indeed dependent on the media compositions (glutamate availability).

Altogether, our results suggested that under a favorable condition (*e.g*. high glutamate concentrations in the media) the probability of premature germination is lower. However, when the condition is not favorable (which results in higher *k*_−2_) or when forespores have morphological errors (which results in lower *k*_1_), forespores are more prone to germinate prematurely.

### Spores produced in media containing ThT are less resistant against wet-heat

Considering that the quality/quantity control of endospores poses a challenge in our society (19), we wondered if the above understanding enables us to propose a novel way of controlling the quality and quantity of endospores.

Ramirez-Guadiana *et al* reported that the most dramatic increase in premature germination was observed with the deletion of *ylbJ* (later renamed as *spoVV*) (18). SpoVV, a concentrative nucleoside transporter, translocates dipicolinic acid (DPA; pyridine-2,6- dicarboxylic acid) against the concentration gradient across the outer forespore membrane from the mother-cell compartment to the intermembrane space (32). Concentrative nucleoside transporters are cation symporters: their driving force is the electrochemical concentration gradient of monovalent cations (typically sodium) (33), and thus, the cation accumulation on forespore surfaces can provide a driving force for the SpoVV concentrative transporter (Figure 5A). DPA accumulation is an important step in sporulation and is typically associated with the wet-heat resistance of endospores (23). Therefore, the insights from literature and our study make a specific prediction that the endospores prepared with ThT should be less resistant against wet-heat treatments.

**Figure 5.**
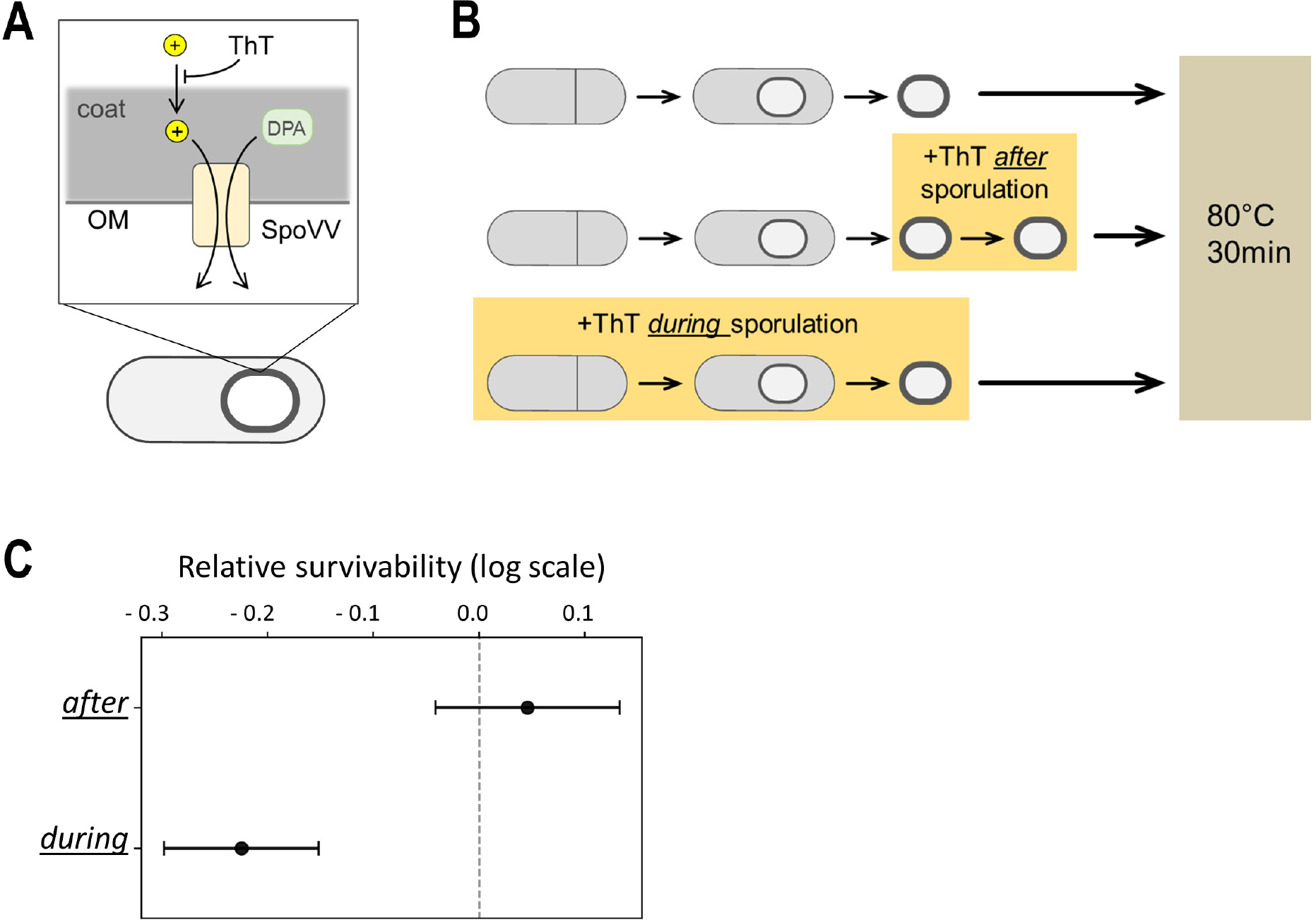
Supplementation of ThT not only decreases the yield of endospores but also decreases the wet-heat resistance level of endospores. A) Diagram illustrating the hypothesis that ThT mimics the phenotype of *ΔspoVV (ylbJ)* strain. SpoVV is a concentrative transporter for dipicolinic acid (DPA), which requires cation concentration gradient for its function. The concentrative transporter SpoVV is located on forespore outer membrane (OM). B) Diagram describing the wet-heat resistance assay shown in panel C. Cells were committed to sporulation (culture for 3.5 hours in RM), then moved to RM with and without ThT. Spores were treated at 80 °C for 30 minutes. See *Methods* for details. C) Survivability of endospores relative to spores produced without ThT (the top one in panel B). Wet-heat survivability is lower when spores are formed with the presence of ThT. Survivability is comparable when ThT is added to the endospores already prepared in RM without ThT. Error bars are 95 % confidence intervals for Poisson distribution based on the number of spores observed.

Following this prediction, we prepared endospores using the sporulation media RM with and without 10 μM ThT, and conducted the wet-heat resistance assays at 80 °C for 30 minutes (Figure 5B). As a control, we also treated endospores with ThT after completion of spore formation (Figure 5B; middle). The survivability of endospores was measured for each sample and normalized to the survivability of endospores prepared without ThT (Figure 5B; top). The endospores exposed to ThT after completing sporulation (Figure 5B; middle) exhibit the survivability comparable to the endospores without ThT (Figure 5C). However, the endospores prepared with ThT are more sensitive to the wet-heat treatment than the endospores prepared without ThT (Figure 5C). This result suggests that supplementation of ThT not only reduces the yield of endospores (Figure 4B) but also diminishes the wet-heat resistance property (Figures 5B and 5C).

## DISCUSSION

Premature germination mediated by GerA was recently reported as a type of quality-control system, however, it was left unclear how this process can be responsive to diverse morphological errors of forespores. By measuring the single-cell dynamics of cations during late sporulation, we demonstrated that forespores poorly accumulating cations preferentially germinate while still inside the mother cells. Our results revealed that the premature germination process is coupled with the external and internal conditions during late sporulation through the cation accumulations on forespore surfaces. This mechanism could add plasticity to the genetically regulated processes of sporulation.

### ThT as a potential chemical to prevent the formation of resistant endospores

We showed that ThT decreases the yield of endospores by promoting the premature germination, while at the same time, lowering the wet-heat resistance of endospores. These results propose a novel usage of ThT as an agent to limit the formation of resistant endospores. We believe this is an attractive possibility since ThT is non-toxic to humans and unlikely to cause undesired side effects; in fact, ThT has been used as a fabric dye and a pen ink. Intriguingly, ThT at 50 μM has been shown to prevent the disruption of muscle sarcomeres during the aging and extend the median lifespan of *C. elegans* by 43~78 % (34). Coincidentally, 50 μM is the ThT concentration at which we observed almost complete abolishment of phase-bright endospore formation. Therefore, ThT at the concentration around 50 μM may bring multiple benefits to the industries where endospore formation poses problems. We also note that ThT is a relatively inexpensive chemical; a liter of 50 μM ThT solution would cost approximately 0.05 US dollars. If ThT alters the yield and quality of other medically important spore-forming bacteria, such as *B. anthracis, B. cereus*, and *C. difficile*, it may offer benefits in the medical sectors as well. Spores of these species are also negatively charged on their surfaces (28, 29), which suggest the mechanism may be conserved among those species. As such, systematic investigation of the impact of ThT in various spore-forming bacterial species would be an important avenue of further research.

### Cation accumulation as an integrative indicator for various internal and external environments

A universal challenge of quality-control systems is to integratively monitor the diverse internal and environmental factors which may be influential for the offspring survival. This is a challenging task to be achieved solely by specific sensor proteins because it requires a wide variety of sensing proteins and integration of inputs. In the case of *B. subtilis* sporulation, our data suggested that the cation accumulation can be seen as a monitoring system affected by both internal and external factors. This mechanism presents an elegant solution to the challenge of sensing diverse environmental factors and morphological errors. An important following research project is to systematically analyze if this mechanism is conserved among different sporangia species.

### Potential roles of premature germination in biofilm colonies

Our study presents a new perspective in the emerging research field of bacterial biofilm electrophysiology (1). We showed the frequency of premature germination depends on the mother cells’ ability to uptake cations from their environment. Because the efficiency of cation uptake can be altered by the electrical signaling in biofilms (2), electrical signaling should, in theory, alter the frequency of premature germination. Intriguingly, in *B. subtilis* biofilms, sporulation is regulated both in space and time; forms a fruiting-body like structure (35). Recent studies suggest the spatio-temporal organization may be regulated by different expression patterns of KinA-E kinases (36), however, the mechanism behind this spatio-temporal organization is still not fully understood. Based on the insight we gained in this study, we suspect that the spatio-temporal organization of sporulation during biofilm formation may also be regulated by electrical signaling and premature germination. Given that premature germination has been seen as a type of programmed cell death (18, 37), it may also provide some benefits to the surrounding cells by providing nutrients or physical space. Such interactions would be pronounced when cells are structurally organized, such as in biofilms (38, 39). Hence, it is plausible to speculate that the cell death through premature germination may provide more significant population-level impacts in biofilms. In our future research, we shall determine the potential roles of biofilm electrical signaling to sporulation, and *vice versa*.

## ACKNOWLEDGEMENTS

We thank GM Suel, P Schafer, DY Lee, T Qagatay, A Kuchina, A Saggese, and the members of the Asally lab (J Stratford, M Delise, I Lopez-Grobas, and C Edwards) for their comments to the drafts of the manuscript, E Ricca, L Baccigalupi and R Isticato for generously providing bacterial strains. This study was supported by the start-up fund from University of Warwick, SLS Pump priming fund, and the Royal Society Research Grant to MA, and BBSRC/EPSRC grant to Warwick Integrative Synthetic Biology Centre (BB/M017982/1). PB was supported by a scholarship from the Region Auvergne-Rhone-Alpes and the Université Grenoble Alpes.

---

**Table S1.** Bacterial strains and primers used in this study.

| ***B. subtilis* strains** |  |  |
| --- | --- | --- |
| **Strain** | **Genotype** | **Source** |
| PY79 | Wildtype | Lab collection |
| PY79<br>$\Delta gerA$ | *gerA::spec<sup>R</sup>* | This study |
| PY79<br>$\Delta SpoIVCA$ | *spoIVCA* | (40) |
| PY79<br>$\Delta GerE$ | *gerE36* | (41) |
| PY79<br>$\Delta CotE$ | *cotE::cm<sup>R</sup>* | (42) |
| PY79<br>$\Delta CotB$ | *cotB::spc<sup>R</sup>(insertion)* | DB068 (43) |
| PY79<br>$\Delta CotG$ | *cotGH::neo<sup>R</sup>;*<br>*amyE::cotGstopcotH* | (44) |
| PY79<br>$\Delta CotBG$ | *\Delta cotG\Delta cotH::neo<sup>R</sup>;*<br>*amyE::cotGstopcotH* | Gift from Loredana Baccigalupi, University of Naples Federico II |
| PY79<br>$\Delta CotC$ | *cotC::spec<sup>R</sup> (insertion)* | (45) |
| PY79<br>$\Delta Cout$ | *cotU::neo<sup>R</sup> (insertion)* | (45) |
| ***E. coli* strains** |  |  |

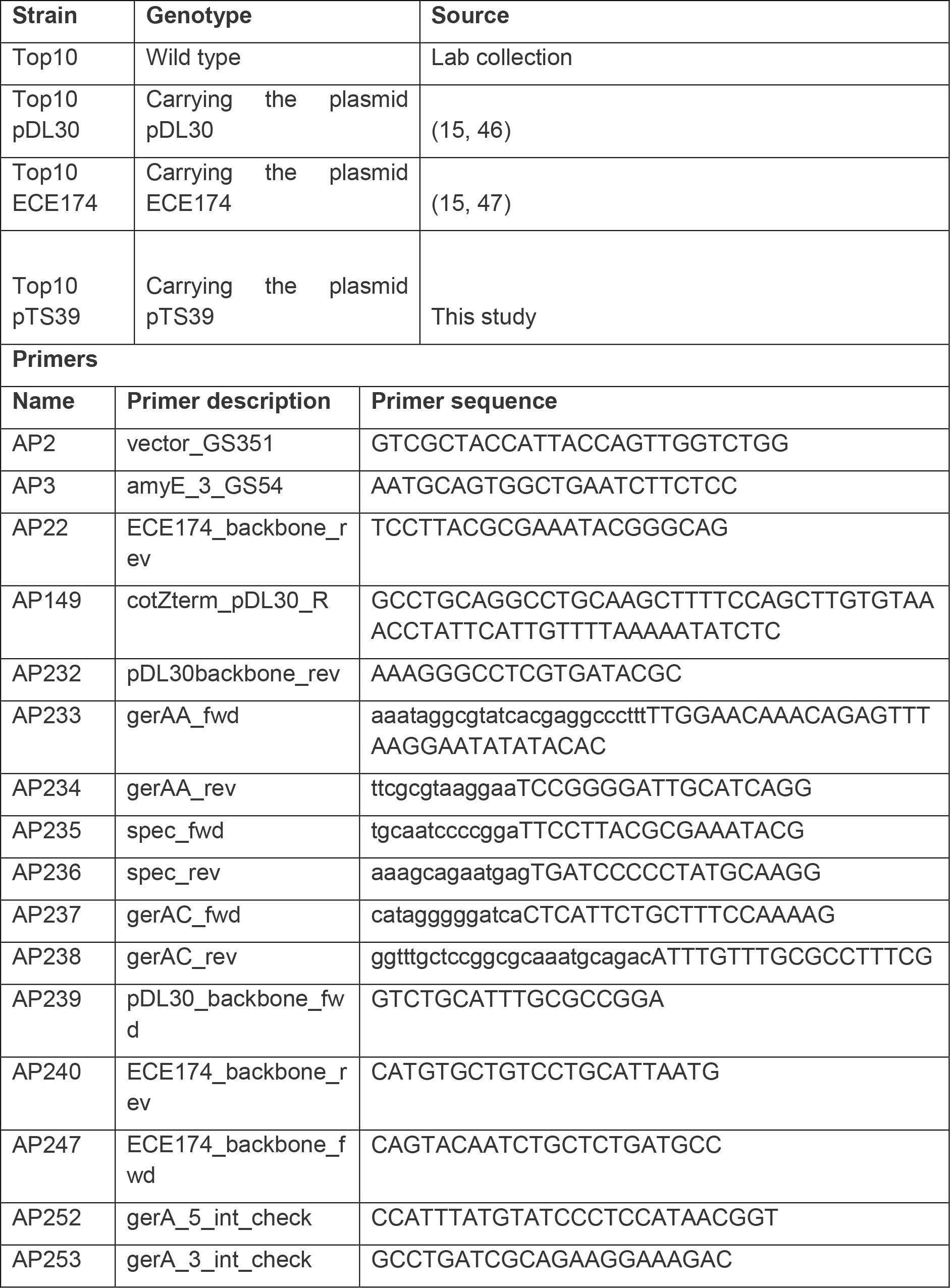

## MATERIALS AND METHODS

### Growth Conditions of *B. subtilis* and *E. coli*

The strains of *B. subtilis* and *E. coli* used in this study are listed in Table. All strains were routinely grown in Lysogeny Broth (LB) or LB agar plates at 37 °C. When culturing the strains carrying antibiotic-resistance genes, appropriate antibiotics were supplemented in LB at the concentrations of 100 μg/mL ampicillin, 300 μg/mL spectinomycin, or 5 μg/mL chloramphenicol.

### Strain constructions

All strains and primers used in this study are listed in the Table S1. PY79 Δ*gerA* strain was derived by transforming pTS39 plasmid using the standard one-step *B. subtilis* transformation procedure described previously (48). For the construction of pTS39 plasmid, the upstream and downstream regions of *gerAA* gene locus are amplified by Polymerase Chain Reaction (PCR) with PrimeSTAR Max DNA Polymerase (Takara) using the primer sets AP233/AP234 and AP237/AP238, respectively. The sequences of these primers, as well as all the other primers used in this study, are listed in Table. The plasmid backbone and spectinomycin-resistance cassette were amplified from pDL30 plasmid (15) with the primer sets AP232/AP239 and AP235/AP236. The PCR products were analyzed by electrophoresis and subsequently purified from agarose gel using gel extraction kit (QIAGEN). The purified DNA fragments were then assembled using the Gibson assembly kit (NEB) and transformed into chemically competent *E. coli* cells (Top10) using the standard heat-shock transformation method. Cells were then plated onto LB agar plates with ampicillin and cultured overnight at 37°C. The colonies obtained were screened by colony PCR using the primer sets AP233/AP238. The plasmids were then extracted from the positive clones and sent for the Sanger sequencing service by the Source BioScience, plc. The resultant sequence data were aligned to the sequence assembly *in silico* using Benchling (benchling.com). Once the sequence is confirmed, pTS39 plasmid was linearized with the restriction enzyme ScaI, and transformed into wildtype PY79. The colonies grown on LB agar plate with spectinomycin were screened by colony PCR using the primer sets AP252/AP149 and AP253/AP3 to check the double crossover homologous recombination. The positive clones were stored in 25 % glycerol at −80 °C.

### Time-lapse microscopy of sporulation with Resuspension medium (RM)

The sporulation dynamics of *B. subtilis* cells were observed as previously described (15), using the fluorescence microscopy Leica DMi8 equipped with an automated stage, Hamamatsu Orca-flash 4.0 scientific CMOS (complementary metal-oxide-semiconductor) camera, and a PeCon incubation system. The objective lens HCX PL FLUOTAR 100x/1.30 OIL PH3 was used for all microscopy assays. For sporulation time-lapse microscopy, overnight LB cultures of *B. subtilis* were resuspended in 20 % (v/v) LB at OD_600_ ~ 0.1 and incubated at 37 °C with shaking until reaching OD_600_ 0.6 - 0.8. The cultures were then centrifuged at 4,000 rpm (revolutions per minute) for 10 minutes, and resuspended in equal volume of pre-warmed Resuspension Medium (49) (RM; composition per 1 liter: 46 μg FeCl2, 4.8 g MgSO4, 12.6 mg MnCl2, 535 mg NH4Cl, 106 mg Na2SO4, 68 mg KH2PO4, 96.5 mg NH4NO3, 219 mg CaCl2, 2 g L-glutamate), and then incubated for 3hr at 37 °C in a shaking incubator prior to microscopy assay. The cultures were then deposited on LMP (Low Melting Point) agarose pads prepared as described previously (15). Briefly, 1.8% (weight/volume) LMP agarose was dissolved in RM without glutamate by microwave and left to cool down before adding glutamate. When specified, fluorescent dyes or inhibitors are supplemented to the RM at the following concentrations; 1 μM APG-2 AM (TefLab), 1 μM Fluo-4FF AM (Molecular Probes), 10 μM (unless specified otherwise in the figure legend) Thioflavin T (Sigma-Aldrich), 10μM valinomycin (Sigma-Aldrich), 20 nM TMRM (Molecular Probes). 1 mL of the RM-LMP agarose solution was placed onto a cover glass (22 millimeters by 22 millimeters) and covered by another cover glass of the same size. Once agarose is polymerized, the coverslip on the top was removed, and 2 μL of cell cultures were deposited onto the agarose pads. The pads were cut and placed in a glass-bottom dish (Willco, HBST-5040) to be used for microscopy assays.

### Mathematical model

We model the dynamics of the cytoplasmic and forespore-bound cations with the following coupled differential equations:

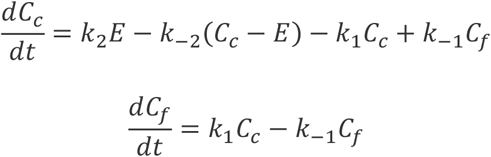

where *C_c_* is the concentration of cytoplasmic cations and *C_f_* is the concentration of cations bound to forespore. We define the following reactions rates:

*k*_1_: association constant of the cytoplasmic cations to the forespore
*k*_−1_: association constant of the cytoplasmic cations to the forespore
*k*_2_: import rate of the cation pump
*k*_−2_: diffusion rate of the cation channel

Considering the development of forespore and increase with time in its negative charge, we assume that *k*_1_ grows monotonically with time. We assume this time dependence as:

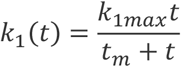

This equation implies that *k*_1_ grows and eventually saturates to a value *k*_1*max*_, and *t_m_* represents the time at which the crossover from linear to zero-order behavior occurs. The numerical simulation was performed using Python, and the parameters used in the simulations are specified in the figure legend.

### Image analysis

The time-lapse microscopy images were analyzed with Fiji/ImageJ (National Institutes of Health) (50). The ROI (region of interest) manager function of ImageJ was used to register the fore-/endo-spore regions of sporulating cells on each frame of time-lapse microscopy images.

The centroid coordinates and pixel intensity values were recorded for each ROI. The coordinate was used to track individual cells using Trackpy (github.com/soft-matter/trackpy), a Python package for particle tracking. For the quantification of ThT fluorescence on endospores, a bespoke imageJ macro was developed to detect spores. For the quantification of premature germination, phase-bright and phase-dark spores were counted on the frame 30 (600 minutes into time-lapse microscopy experiment).

### Quantitative analysis

Although making binary decisions of analyzed results using p-values is still common in biological science, the use of null hypothesis significance testing (NHST) approach for this purpose has been seriously questioned. It has been demonstrated that NHST is often unreliable with the common experimental settings in biological science (51, 52). It has also been argued that misuse of NHST and p-values may be an underlining reason for the challenges in irreproducible research. 95 % Confidence Interval (CI) has been proposed to use as an alternative approach to statistical interpretation. As such, in this study, we show the 95% confidence intervals whenever appropriate to do so. For the frequency of premature germination, we calculated the confidence intervals based on the number of events observed. The calculation was made by using the stats package in scipy (SciPy.org). Numpy (NumPy.org) and Pandas (Pandas.PyData.org) were used to calculate the standard statistical values (mean, standard deviations). All plots were created using the Python packages, matplotlib (github.com/matplotlib/matplotlib) and seaborn (github.com/mwaskom/seaborn).

### Spore preparation

Sporulation of *B. subtilis* cells was induced by the resuspension method of Sterlini & Mandelstam as described previously (49). A single colony of *B. subtilis* was inoculated in 5 mL LB and incubated for an overnight at 37°C in a shaking incubator (200 rpm). The culture was then diluted to OD_600_ of 0.1 and incubated at 37°C in prewarmed 20% (volume/volume) LB for 3 hours to reach OD_600_ of 0.6 - 0.8. The culture was centrifuged for 10 minutes at 4,000 rpm at room temperature (RT) and resuspended in an equal volume of pre-warmed RM. After an overnight incubation in RM at 37°C in a shaking incubator, the culture was centrifuged for 10 minutes at 4,000 rpm and washed 3 times in distilled water. To remove the vegetative cells from the solution, the culture was treated with 100 μg/mL lysozyme in 10mM Tris-HCl (pH 6.8) and incubated for an hour at 37°C with aeration (200 rpm). The resultant solution was subsequently washed in 1M NaCl, milli-Q water, 1M KCl and 10 times in milli-Q water. All the centrifugation steps for washing were performed for 10 minutes at 4,000 rpm at RT. To ensure endospores are purified, the resultant solutions were inspected under microscope using 1.8% (w/v) low-melting-point agarose pad containing PBS.

### Effects of ThT to sporulation in liquid RM

Liquid sporulating cultures in RM were prepared in the same manner described above in the section for the spore preparation. After 3.5 hours of incubation in RM, the cultures were supplemented with the final concentrations of 0 μM, 1 μM, 10 μM or 50 μM ThT and incubated at 37°C overnight in a shaking incubator (200 rpm). The cultures were then examined under microscope using LMP-agarose pads. The number of phase-bright spores was counted using Fiji/imageJ Macro using the particle analyzer function of imageJ.

### Germination assays

For the germination assay by a spectrophotometer, purified free spores were diluted to OD of ~0.8 in 1 mL sterile milli-Q water with various concentrations of salts as specified in the figure legends. The germination assays were conducted without heat shock treatments commonly used in germination assay. L-alanine was added to the final concentration of 5mM and OD_600_ was monitored using a spectrophotometer (JENWAY 7305) every 10 min for 120 min. The germination efficiencies (%) in various salt conditions were calculated as the percentage drops in OD_600_ as; 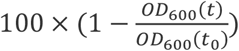. To examine the impacts of salt to germination efficiency, the data of the last time point (*t_120_*)was plotted as a function of salt concentrations.

### Wet-heat resistance assay

The spores were prepared as described above in the spore preparation section. Spore suspensions at OD600 ~1.0 were prepared and incubated at 80 °C for 30 minutes (heat treatment). The samples (heat treated and non-heat treated) were serially diluted and plated onto LB plates (1.5% agar) with three replica plates for each condition and serial dilution. After an overnight incubation in a 37 °C incubator, colonies on LB plates were counted, and the average values of three replica plates were used for analysis. The numbers of colonies appeared with heat-treated samples were divided by the number of colonies appeared with non-heat-treated samples to calculate the survival rate. We also performed time-lapse microscopy assay with the endospores on LB agar pads and quantified the number of endospores that outgrew in 2 hours at 37 °C.

---

**Figure S1.**
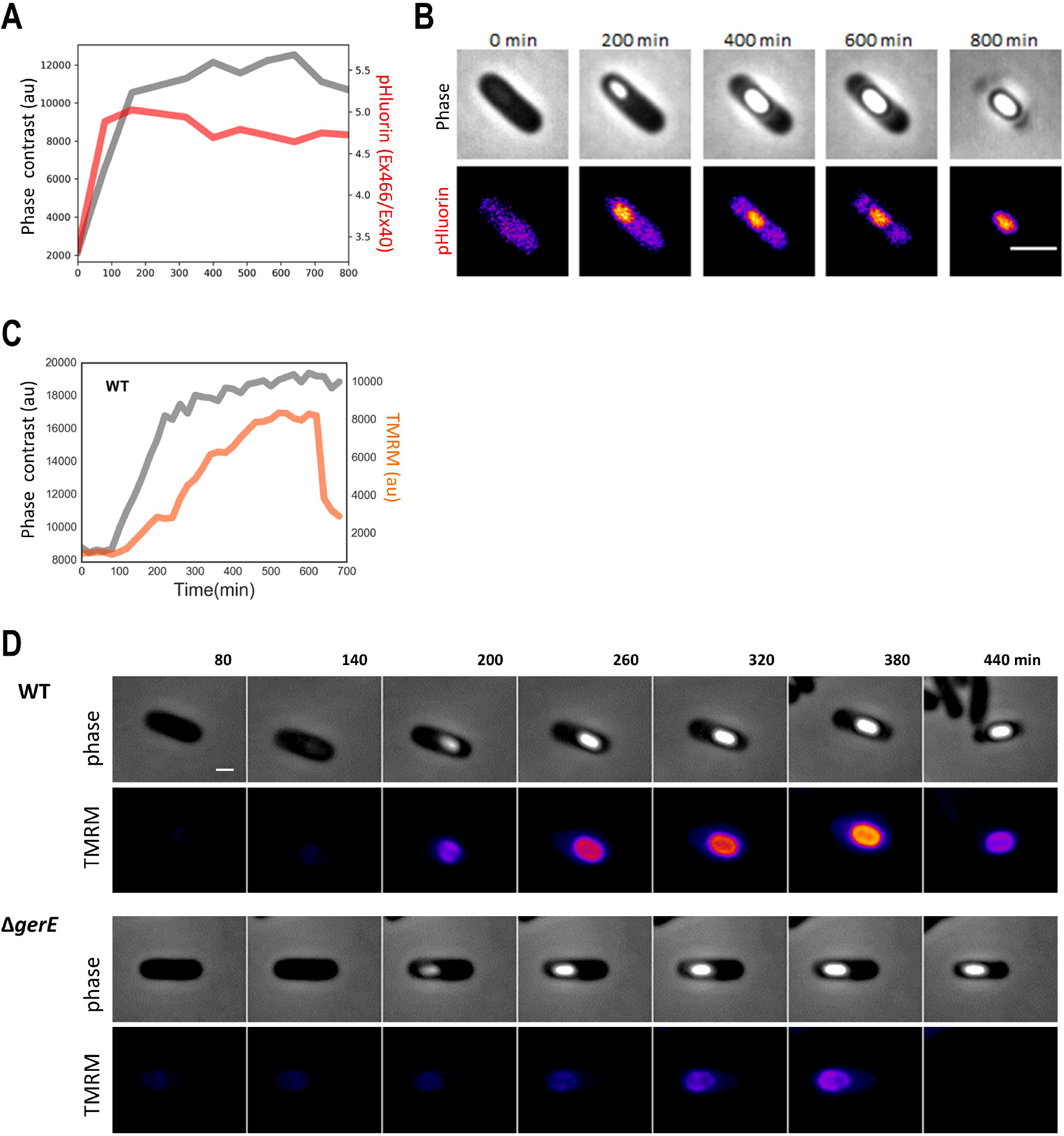
TMRM and pHluorin dynamics during sporulation. A) Time series of forespore/endospore intensities of phase contrast channel in grey and pHluorin in red (calculated as Ex466/ Ex400 ratio) in wild type strain. Time series shows a steep acidification of forespore cytoplasm coinciding with the occurrence of phase-bright spore. Time series is representative from five independent experiments. B) Film-strip images showing the phase-contrast channel (upper) and pHluorin fluorescence channel (lower) of a live single-cell sporulating cell in wild type strain. The scale bar is 2 μm. C) Time series of forespore/endospore intensities of phase contrast channel (grey) and TMRM fluorescence (orange) in wild type strain. Time series shows a gradual increase of TMRM, which drops upon mother-cell lysis. Time series is representative from five independent experiments. D) Film-strip images showing the phase-contrast channel (upper) and TMRM fluorescence channel (lower) of a live single-cell sporulating cell for wild type and *gerE* deletion mutant. The scale bar is 1 μm. Color scale is shown in the left-end panel for TMRM.

**Figure S2.**
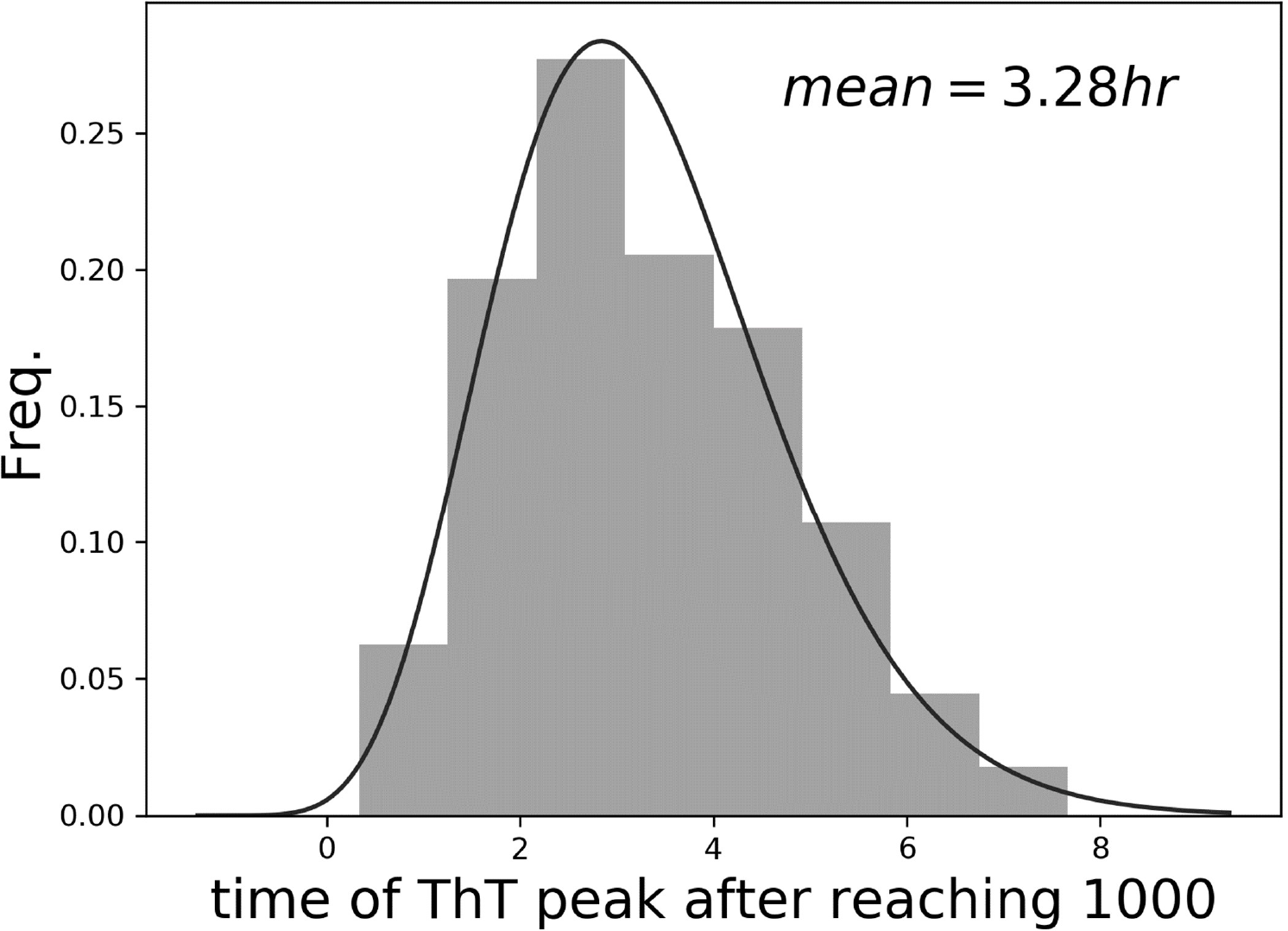
Heterogeneity in the duration of late sporulation. Histogram of the duration of which it takes to mother-cell lysis when time series are aligned as presented in Figure IE. Same dataset was used as Figure 1D and 1E.

**Figure S3.**
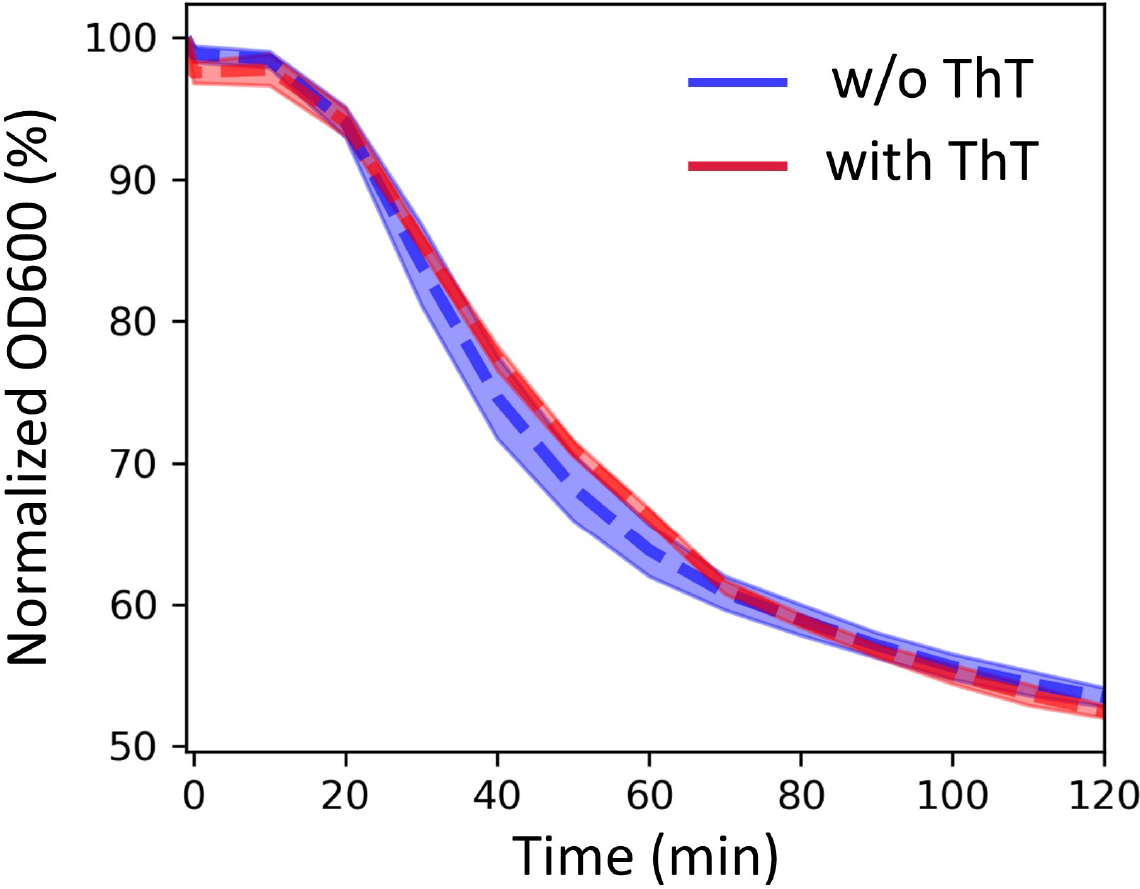
ThT influences premature germination during spore formation but has no effect on germination in purified mature spores. ThT has no effect on germination of mature purified spores. Herein, we compare the germination profile of purified spores and purified spores stained with ThT. Error bars are standard deviations.

**Figure S4.**
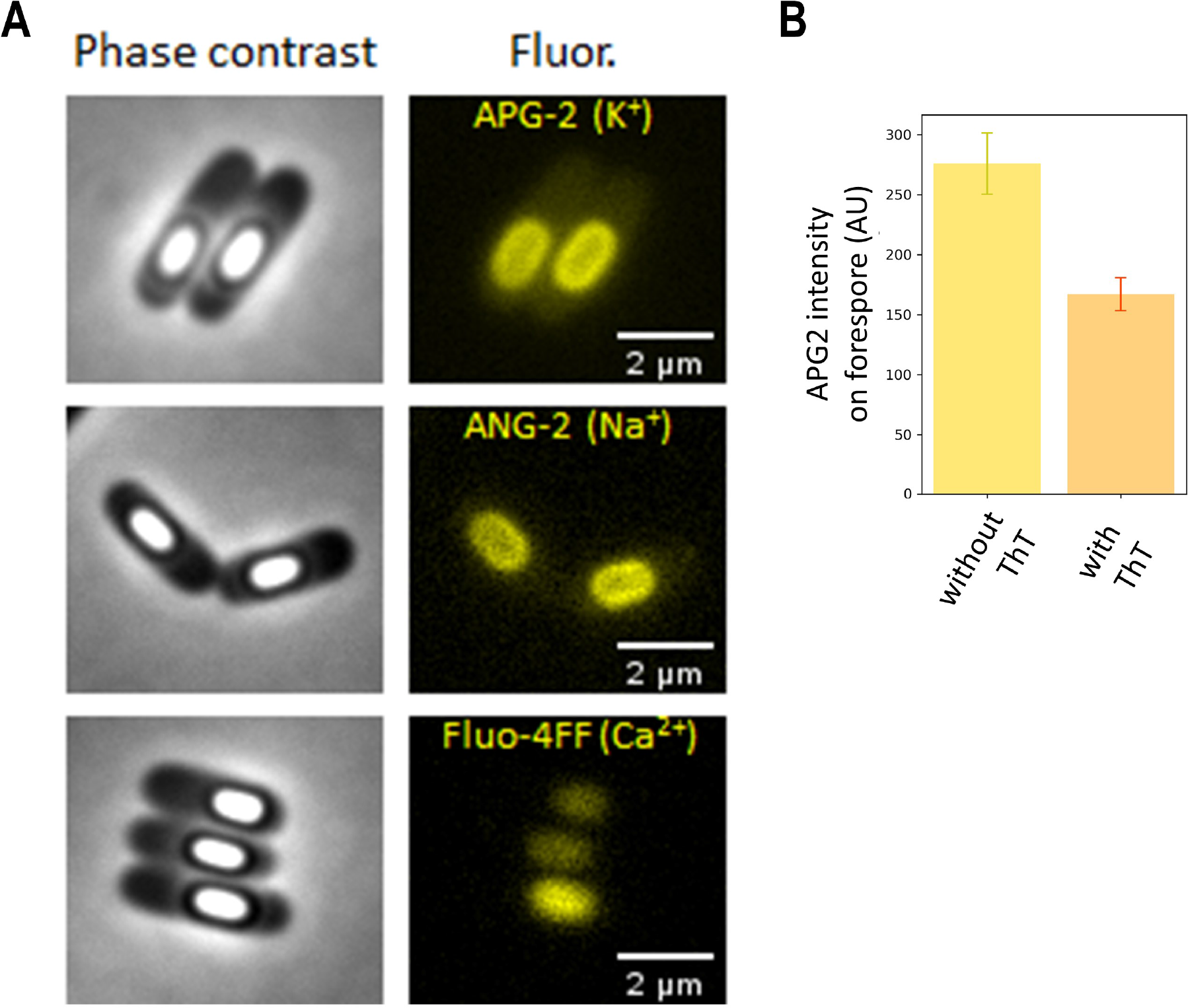
Potassium and sodium localise in spore protective layers while calcium localises in the spore core. A) Positively charged ions localise in different spore compartments. Calcium in the spore core, potassium and sodium in protective layers surrounding the spore. Microscopy images show phase contrast (left, grey), and APG2-AM, ANG2-AM and Fluo-4FF fluorescence (right, yellow) on spores. Scale bar, 2 μm. B) Quantification of APG2-AM fluorescence on forespore without and with ThT in the media. 30 sporulating cells were analysed for each condition. Error bars are 95% confidence intervals for the Normal distribution.

**Figure S5.**
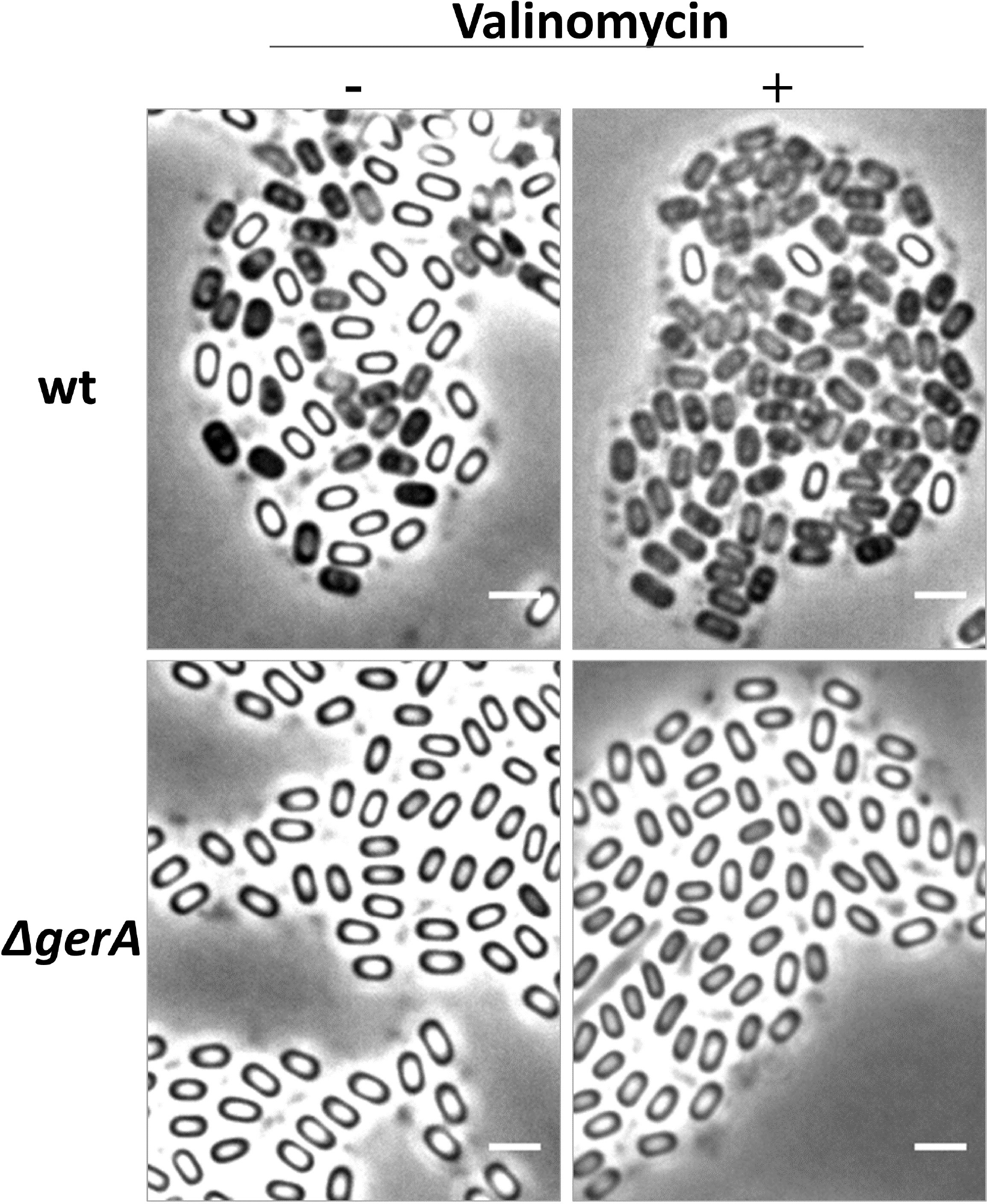
Valinomycin affects sporulation of wild type strain but not of *gerA* mutant. Microscopy images showing representative samples of wild type and *gerA* mutant strain. Phase contrast (grey) images show the proportion of white and black spores without and with valinomycin treatment. Scale bar, 2 μm.

## Supplementary Movies

### Movie 1 Dynamics of ThT during sporulation

Fluorescence time-lapse microscopy images corresponding to main Figure 1 B and C.

### Movie 2 pHluorin dynamics during sporulation

Time-lapse microscopy images showing a sporulating cell in phase contrast (left) and pHluorin dynamics (right). The intensity in the pHluorin is the ratio Ex466/Ex400. Lighter colours indicate low pH and darker colours indicate high pH.

### Movie 3 TMRM dynamics during sporulation

Movie xz. Time-lapse microscopy showing a sporulating cell in phase contrast (left) and TMRM fluorescence (right).

### Movie 4 Phase contrast images of live widtype cells during sporulation

Time-lapse microscopy showing sporulating wildtype cells in phase contrast. Premature germination is evident as developing forespores turn phase-dark in the mother cell. Scale bar, 2 μm.

### Movie 5 Phase contrast images of live Δ*gerA* cells during sporulation

Time-lapse microscopy showing *gerA* sporulating cells in phase contrast. No premature germination are detected with this mutant strain. Scale bar, 2 μm.

